# Mechanical vibration modulates regional cerebral blood flow and biomechanical co-variance network in a frequency-dependent manner

**DOI:** 10.1101/2022.06.28.498036

**Authors:** Linghan Kong, Suhao Qiu, Yu Chen, Zhao He, Peiyu Huang, Qiang He, Ru-Yuan Zhang, Xi-Qiao Feng, Linhong Deng, Yao Li, Fuhua Yan, Guang-Zhong Yang, Yuan Feng

**Author notes:** Contributed Equally (Linghan Kong, Suhao Qiu). Address for correspondence: Guang-Zhong Yang, Ph.D., Yuan Feng, Ph.D., School of Biomedical Engineering Shanghai Jiao Tong University Shanghai, China.

## Abstract

Human brain experiences vibration of certain frequency during various physical activities such as vehicle transportation and machine operation or accidents, which may cause traumatic brain injury or other brain diseases. However, little is known about what happened to brain after vibration stimuli. Here, with a custom-built electromagnetic actuator, vibration was induced in the brain while cerebral blood flow (CBF) and brain stiffness were measured at 20, 30, 40 Hz for 52 healthy volunteers. With increasing frequency, multiple regions of the brain showed increasingly reduced CBF, while the size of such regions also expanded. The vibration-induced CBF reduction regions largely fell inside the brain’s default mode network (DMN), with about 58 or 46 % overlap at 30 or 40 Hz, respectively. By establishing a biomechanical co-variance network based on tissue stiffness, analysis of small-world properties and modularity showed an increased disruption of the network with increased frequency. These findings demonstrate frequency-dependent features of vibration modulation to brain. Furthermore, the overlap between CBF reduction regions and DMN, and the vibration-induced decrease of biomechanical network connections suggest a interweaved relationship between blood flow, tissue stiffness, and cognitive functions. These may provide critical insights into the mechanical stimulus to brain and vibration-induced brain pathologies.

## 1. Introduction

Vibration is a common mechanical stimulus to our brain during daily life and work, such as in-vehicle transportation and machine operation, which has been strongly associated with symptoms of motion sickness (*1–5*), vibration syndrome (*6*), vertigo, and dizziness (*7*). It is reported driving fatigue, contributing to 20% of automobile crashes, is actually brain dysfunction and impairment resulting from motor vehicle induced brain vibration (*8*). In more severe cases, both short-period dynamic high impact and long-period repetitive moderate vibration can cause traumatic brain injury (TBI) (*9–20*). The etiology of vibration-induced TBI, however, is not yet fully established due to a lack of detailed knowledge about how vibration affects the human brain.

In studies concerning vibration-induced tissue damage in hand-arm vibration syndrome (HAVS), it has been widely observed that vibration causes an early effect of vasoconstriction that leads to obstruction of blood flow and oxygen transportation to the tissue (*21–26*). And the vasoconstriction has also been found to be dependent on both the magnitude and frequency of the vibration (*27, 28*). On the other hand, patients with TBI also show reduced cerebral blood flow (CBF), a major physiological marker of brain functions, in the bilateral frontotemporal regions, together with structural changes of the cerebral blood vessels and disintegrity of white matter (*29–31*). Widespread deficits of CBF have also been found in grey matter in correlation with severity of TBI and cognitive dysfunction(*32*).

Experimental and computational studies appeared to indicate that vibration-induced effects on brain may well be frequency-dependent. Yan et al. (2015) used and animal model to show that significant damage could be introduced to brain after certain periods of vibration at 30 Hz (*33*). Laksari et al. (2018) simulated the motion and strain based on dynamics data from head impacts and found that the brain was more susceptible to vibration at frequencies ranging between 20-40 Hz, with strain peaking at around 30 Hz (*34*). Although animal and simulation studies provided crucial information of the frequency properties of brain, it remains challenging to experimentally characterize the *in vivo* frequency-dependence of vibration-induced effects on brain functions.

Using magnetic resonance elastography (MRE), studies have shown stiffness of the brain increased with frequency monotonically (*35*). In addition, variation of the regional stiffness has also shown to be indicative of brain function (*36*). Although intensive studies have been carried out by analyzing the co-variance network using the structural information, i.e. cortical thickness between brain regions (*37*), little is known about the co-variance network based on the biomechanical information, and its frequency-dependent properties. To explore the frequency-dependent behavior of the brain, we built a bespoke electromagnetic actuator to be worn inside the MR head coil to apply mechanical vibration to the brain. CBF and brain tissue stiffness were acquired at precise frequencies of 20, 30, and 40 Hz, respectively. The frequency-dependent changes of CBF and biomechanical co-variance network were analyzed. These results could help discover the frequency-dependent properties of brain tissue from physiological and biomechanical perspectives.

## 2. Results and Discussions

### 2.1 Vibration decreased CBF with increasing frequency

To investigate the effect of vibration on CBF, we conducted two sessions of measurement (**Figure 1b)**, namely Session 1 (resting-state session) and Session 2 (vibration session). During Session 1, brain anatomy (T1W images) and CBF (3D-ASL) in the absence of vibration were acquired as references. During Session 2, brain stiffness during vibration (Vibration/MRE) and CBF immediately after vibration (3D-ASL) at frequencies of 20, 30, 40 (Hz) were acquired twice. These frequencies were chosen because of their significant role in inducing TBI (*34*).

**Figure 1.**
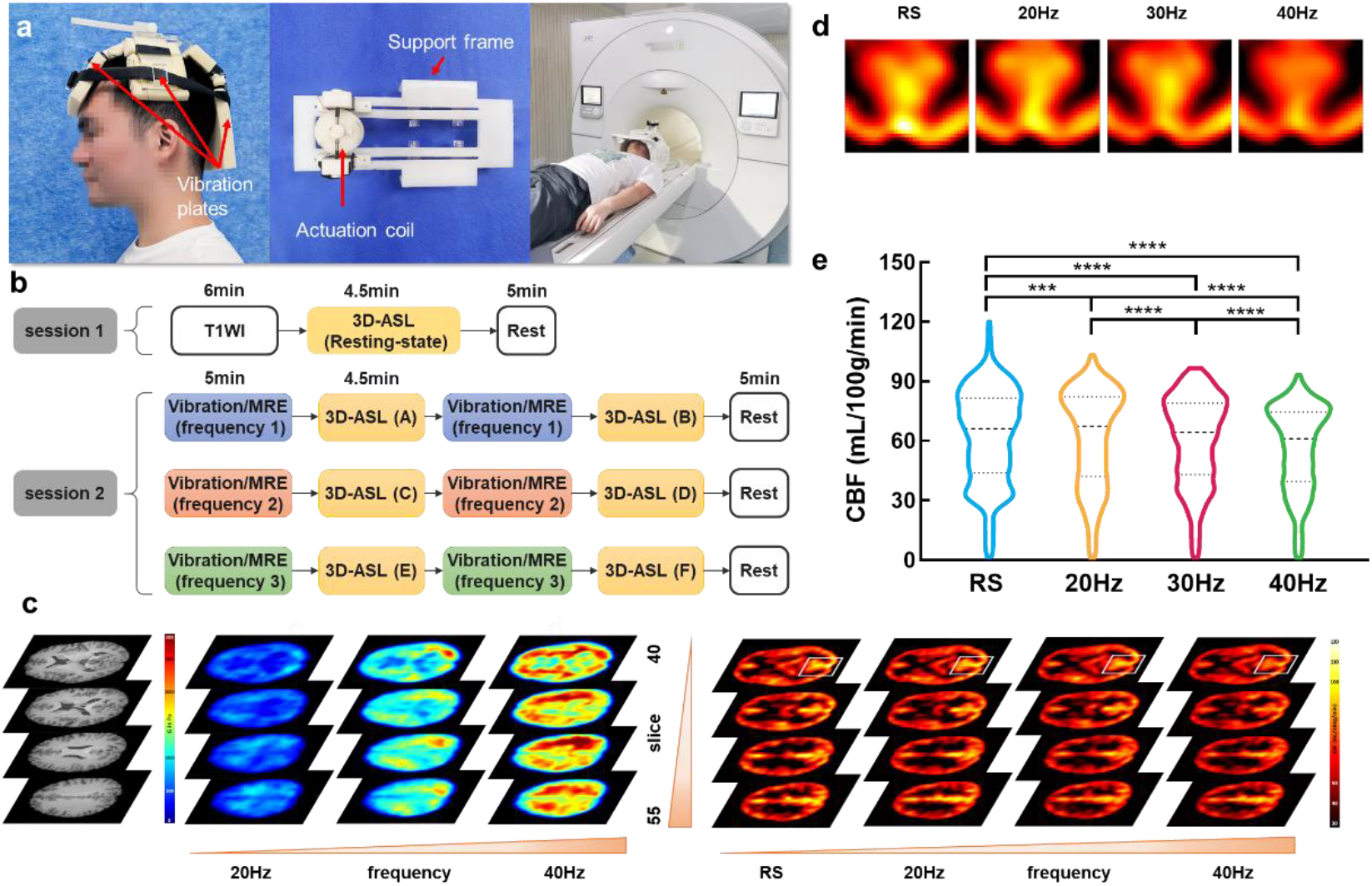
Experimental set-up and procedure for measuring the vibration effect on brain. (a) Custom-built, head worn electromagnetic vibration actuator that transmitted vibration to the brain. The vibration plates were attached with soft sponge pads to ensure the scanning was comfortable. (b) The imaging protocol includes two sessions. Anatomical images (T1W) and CBF maps in the absence of vibration were acquired as references in session 1. Shear stiffness and CBF maps were acquired in the presence of vibration at 20, 30, 40 Hz, respectively. The stiffness and CBF measurements were repeated once more to ensure experimental reproducibility. The order of frequency scanning between 20, 30, 40 Hz was randomized to minimize the potential influence of sequential measurement order. The actuator was used for both vibration actuation and MRE acquisition (*38*). A supine position was adopted for all imaging procedures. (c) Typical anatomical images (T1W) and CBF and stiffness maps of the brain acquired from one volunteer subject in the absence (resting state, RS) or presence of vibration for 5 minutes at different frequencies (20Hz, 30Hz, 40Hz, respectively). The white rectangle box in the top row of CBF map shows an example region (PCC and PCUN). (d) Magnified view of the boxed region in the top row of CBF maps in panel a-b, showing decreasing CBF with increasing vibration frequency (RS to 40 Hz from left to right). (e) Statistical analysis results of quantitative CBF data acquired from all subjects, showing significant decrease of CBF after vibration was applied with increasing frequency (RS vs. 20Hz vs. 30 Hz vs. 40Hz, ***:p<0.001; ****:p<0.0001).

Typical anatomical T1W images, CBF maps, and stiffness maps measured at either the resting state (RS) or immediately after vibration at 20, 30, 40 Hz were shown (**Figures 1c**). For each frequency, the measurement of stiffness and CBF were repeated twice to ensure experimental reproducibility. No significant differences were observed in any clusters at the same frequency (paired student t-tests, p>0.05). Therefore, the mean values of the CBF maps at the same frequency from the two repeated measurements were used for analysis. A magnified view of CBF maps at the occipital lobe (**Figure 1d**) showed the intensity appeared to be decreasing with increasing frequency. Statistical analysis results confirmed that CBF in the boxed region decreased significantly in the presence of vibration with increasing frequency (**Figure 1e**, paired student t-tests, RS vs. 20 Hz vs. 30 Hz vs. 40 Hz, p<0.05).

### 2.2 Vibration with increasing frequency induced expansion of the CBF reduction regions

Compared with the resting state, the regions with significant CBF reduction in the brain after exposure to vibration at 20, 30, 40 Hz were analyzed (paired student t-test, p<0.05). As compared with the resting state, vibration at 20 Hz induced no significant difference in CBF in the whole brain (data not shown). However, vibration at 30 Hz versus the resting state induced a significant decrease of CBF in the region of PCC, PCUN, ITG, IPL, ANG, SMG, SFGmed, TPOmid, TPOsup, PreCG, and SFG (p_corr_<0.05, **Figure 2a**). And vibration at 40 Hz versus resting state induced a significantly decrease of CBF in the region of PCC, PCUN, ITG, IPL, ANG, SMG, SFGmed, TPOmid, TPOsup, PreCG, SFG, and MFG (p_corr_<0.05, **Figure 2b**).

**Figure 2.**
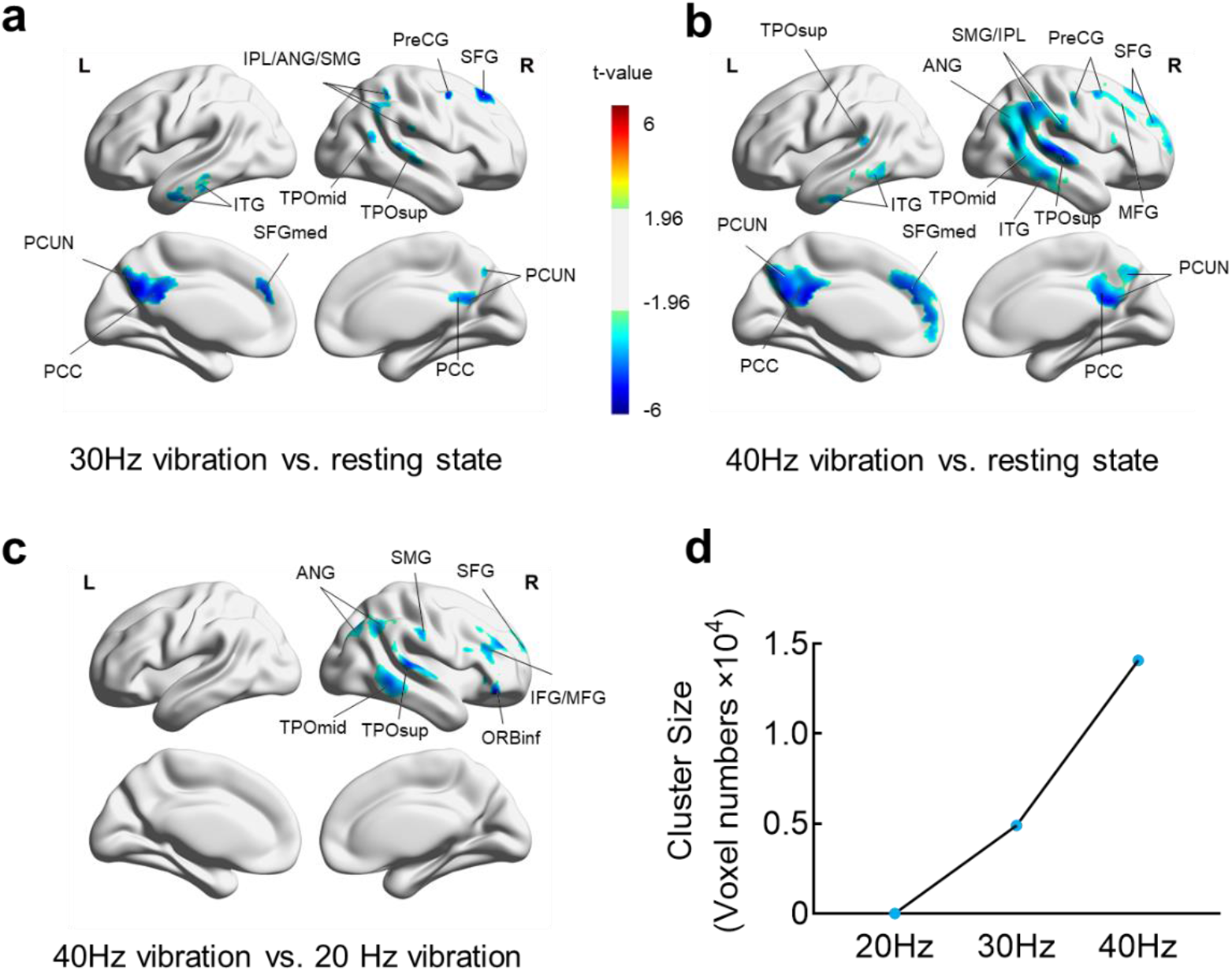
Comparison of the regions with significant CBF reduction in the brain after exposure to vibration at different frequencies. (a) CBF reduction regions of the brain after 30Hz vibration vs. resting state; (b) CBF reduction regions of the brain after 40Hz vibration vs. resting state; (c) CBF reduction regions of the brain after 40Hz vibration vs. 20Hz vibration. (d) Quantified cluster size of regions with significant decrease of CBF versus vibration frequency. Note: Plots were made on the AAL brain atlas. All significant maps were corrected for permutation test and display significant voxels at p<0.05, ANG (angular gyrus), CUN (cuneus), IPL(inferior parietal gyrus), ITG (inferior temporal gyrus), MFG (middle frontal gyrus), ORBinf (inferior frontal gyrus, orbital), PCC (posterior cingulate cortex), PCUN (precuneus), PreCG (precentral), SFGmed (superior frontal gyrus, medial), SMG (supramarginal gyrus), TPOmid (middle temporal gyrus), TPOsup (superior temporal gyrus).

In comparison to CBF maps acquired in the presence of vibration at various frequencies, we found differences of CBF in ROIs between vibrations at 20 Hz and 40 Hz. Specifically, the significant decrease of CBF occurred in the region of ANG, SMG, SFG, TPOmid, TPOsup, IFG, MFG, and ORBinf (p_corr_<0.05, **Figure 2c**). However, no significant difference of CBF was observed between the experimental group with vibration at either 30 Hz versus 40 Hz, or 20 Hz versus 30 Hz. The size of the cluster with significant CBF reduction increased from 0 to about 1.5×10^4^ as the vibration frequency increased from 20 to 40 Hz (**Figure 2d**). Detailed statistics of the cluster reports are given in the **Supplement Information Table S2**.

Compared to the resting state, we observed distinct frequency-dependent regional reduction of CBF, i.e., the region with significant CBF reduction increased with increasing frequency of the vibration. This finding is consistent with that vibration causes an early effect of vasoconstriction in HAVS, which is frequency-dependent(*27, 28*). Studies have shown that CBF can be reduced significantly after brain injury in cases such as mild TBI and sports-related concussion (*39–44*), which is also consistent with our finding of significant CBF reduction after vibration. The method reported in the present study suggests that the vibration-based quantitative measurement technique has the potential to simulate impact and vibration to the human brain while accessing the physiological responses of the brain. In addition, the proposed method also indicated the potential of modulating CBF using vibrations.

### 2.3 Vibration-induced CBF reduction regions overlapped with the brain’s default mode network

To explore how CBF changes could affect brain function, we compared the brain regions showing significant CBF reduction with the 7 brain networks known for cognitive functional connectivity (*45*). The number of overlapping voxels between the regions with significant CBF reduction and those in each of the 7 brain network was estimated for each frequency. We found a large overlap between the regions with significant CBF reduction and the Default Mode Network (DMN) in the brain after vibration at either 30 Hz **(Figure 3a)** or 40 Hz **(Figure 3b)**. Quantitatively, 58% of voxels in the significant region were in DMN, accounting for 9.03% of DMN in the brain after vibration at 30 Hz **(Figure 3c)**, and 45.64% of voxels in the significant CBF reduction region were in DMN, accounting for 20.02% of DMN in the brain after vibration at 40 Hz **(Figure 3d)**. Detailed statistics of the overlapped significant CBF reduction regions are shown in **Supplement Information Table S3**.

**Figure 3.**
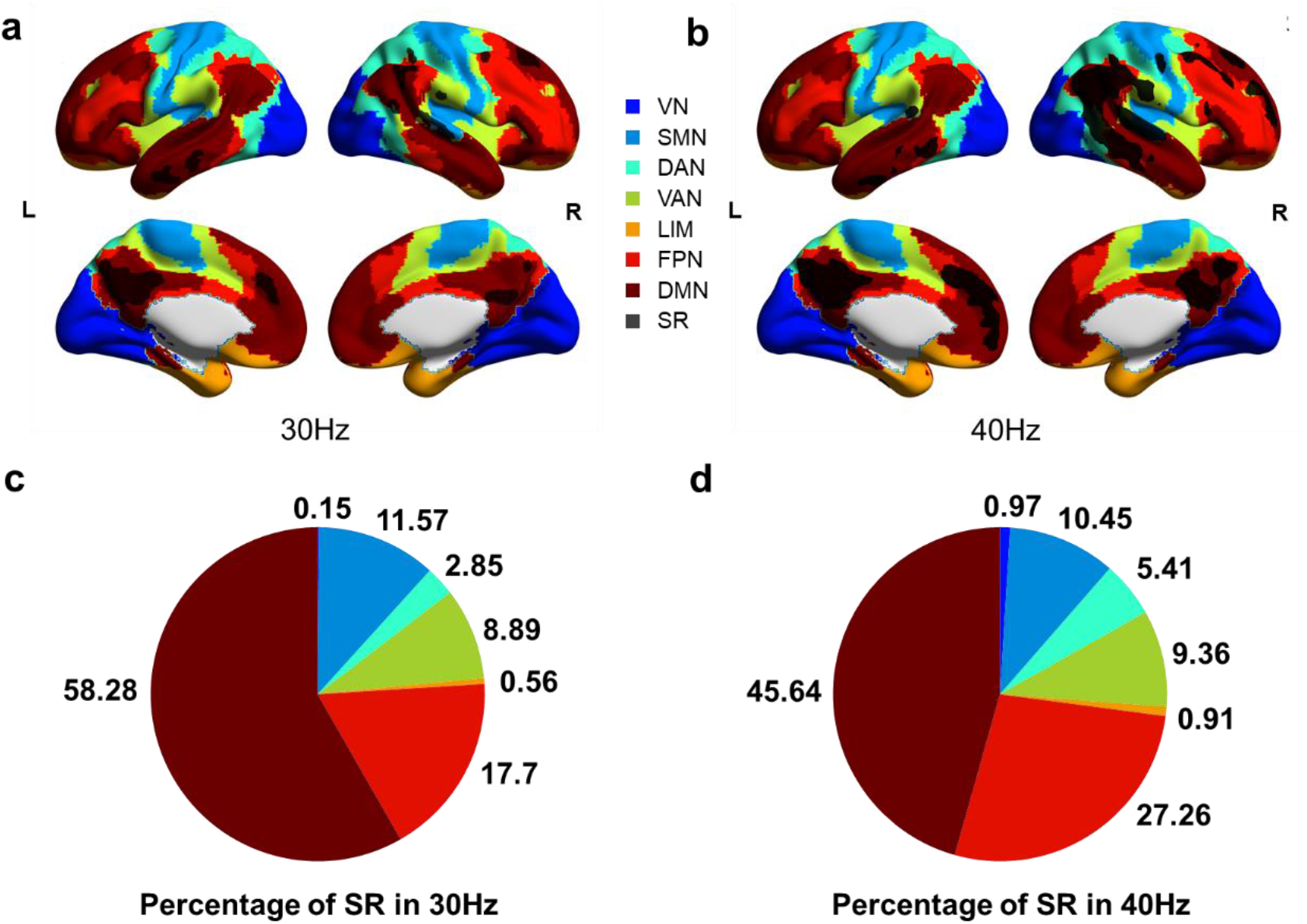
Overlap of the regions with significant reduction of CBF after vibration with the seven brain functional connectivity networks. (a-b) Voxels in regions with significant CBF reduction (SR) after vibration at 30Hz, 40 Hz, respectively, were overlaid with the seven functional connectivity networks of the brain. (c-d) Pie charts of the percentage of voxels in the significant CBF reduction region that overlapped with the seven brain networks after vibration at 30Hz, 40 Hz, respectively. Note: The significant CBF reduction mainly happened within DMN, VN (Visual Network), SMN (Sensory-Motor Network), DAN (Dorsal Attention Network), VAN (Ventral Attention Network), LIM (Limbic), FPN (Frontoparietal Network), DMN (Default Mode Network), SR (Significant CBF Reduction Regions).

In addition, we calculated the mean CBF at each network for all subjects after vibration to the brain at each frequency. Paired student t-tests showed no significant differences of CBF in each network between each vibration frequency **(Supplement Information Figure S1)**, but CBF in the subregions (PCC.L and ACG.R) in DMN was significantly different after vibration at different frequency **(Supplement Information Figure S2)**, possibly due to the average effects of CBF in DMN.

These findings not only prove the concept of characterizing brain functions under specific vibration conditions, but also provide a quantitative measure to understand the relationship between the external mechanical stimulation and the cognitive functions of the human brain. This may be essential for fully understanding the pathophysiological mechanisms of brain damage due to exposure to environmental vibrations as well as potentially help developing better approaches for preventing/treating vibration-induced brain diseases such as TBI. These results are also in line with studies concerning CBF and brain functions of TBI patients. For example, Militana et al. (2016) showed evidence for increased cerebrovascular reactivity and function connectivity in the medial regions of the DMN in college athletes experienced concussion (*46*). Liu et al. (2016) reported that the first 5-mins psychomotor vigilance task decreased the CBF of mild TBI patients in the acute phase in DMN areas (*47*). Therefore, our results bring attention to that vibration as a physical stimulation may induce cognition, memory, or action-related changes, or even injuries.

Our finding of the frequency-dependence of CBF reduction due to vibration has several implications. In the experiments with vibration at frequency of 20, 30, 40 Hz, the size of the regions with significant CBF reduction consecutively expanded when the frequency exceeded 20 Hz. This indicated that in an environment with vibration, absorption or damping for vibration at frequency over 20Hz is desired to protect the brain. Compared with computational studies that showed peak strain in the brain exposed to vibration at 30Hz, our results indicated that during an impact, the brain may respond by changing strain and CBF in different patterns. Moreover, the fact that mechanical vibration could cause CBF reduction in specific regions such as PCC and PCUN in a frequency-dependent fashion may open a new door for using physical modulation of CBF with external vibration as a therapeutic approach to treat certain brain regions.

Another indication of our results is that the vibration may influence the measurements of CBF or other perfusion-related values. Therefore, in terms of measurement order, it is suggested that CBF or BOLD-fMRI tests be carried out before applying any physical stimulation to the brain.

### 2.4 Biomechanical co-variance network

Similar to the co-variance network based on cortical thickness (*48*), the biomechanical co-variance network was constructed (**Figure 4**). The raw biomechanical connection matrixes, FDR corrected (q<0.05) matrices, and the binarized matrices were analyzed (**Figure 4)**. The axis number represented the label of brain region (AAL). The small-world properties of the biomechanical network were analyzed by ANOVA (The small-world properties passed Shapiro-Wilk normality test in all frequency, W=0.9759/0.9191/0.8855 of 20/30/40Hz) with Bonferroni’s multiple comparisons test (**Figure 5**). Significant differences were observed in terms of σ between all 3 vibration frequencies. In addition, a monotonically decreased σ was observed as the frequency increased. Since σ represents a ratio of connection quantities and compactness, this indicates that as the vibration frequency increased, the level of interconnections between brain regions decreased. Studies have shown blood flow is closely related to the tissue stiffness(*49*). Here, as the frequency increased, we observed increased tissue stiffness and decreased CBF. Using a computational model, we demonstrated the coupling between blood flow and tissue stiffness (**Supplementary Figure S3**). The frequency-dependent behavior of brain network further showed interweaved relationship of CBF and tissue stiffness due to vibration modulation.

**Figure 4.**
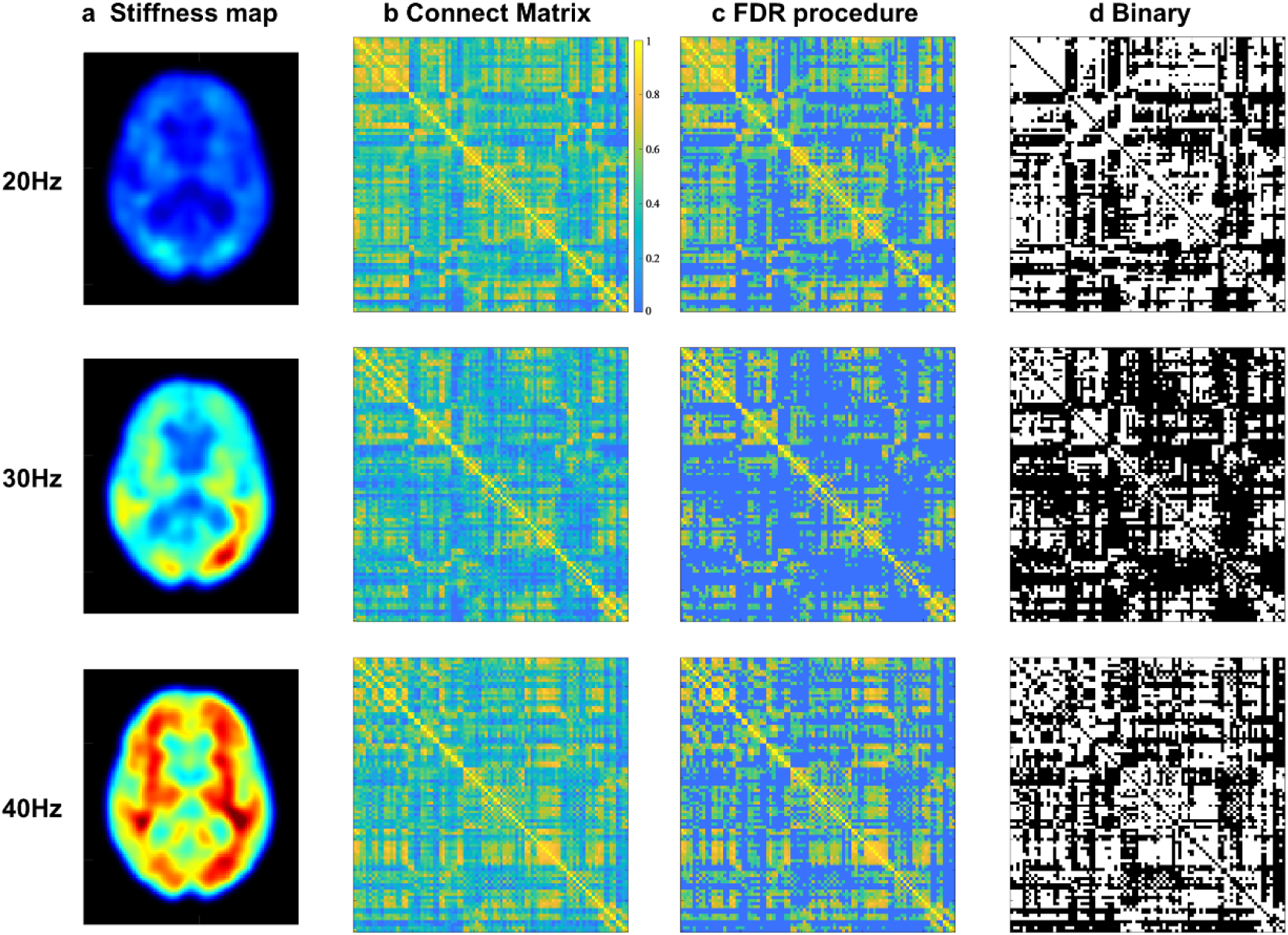
(a) Stiffness maps measured at 20, 30, and 40Hz. (b) The correlation matrix of the cortical stiffness. (c) The correlation matrix after FDR thresholding (q<0.05). (d) The corresponding binary correlation matrix after FDR thresholding. The horizontal and vertical axes represent 90 cortical ROIs using AAL90 parcellation atlas.

**Figure 5.**
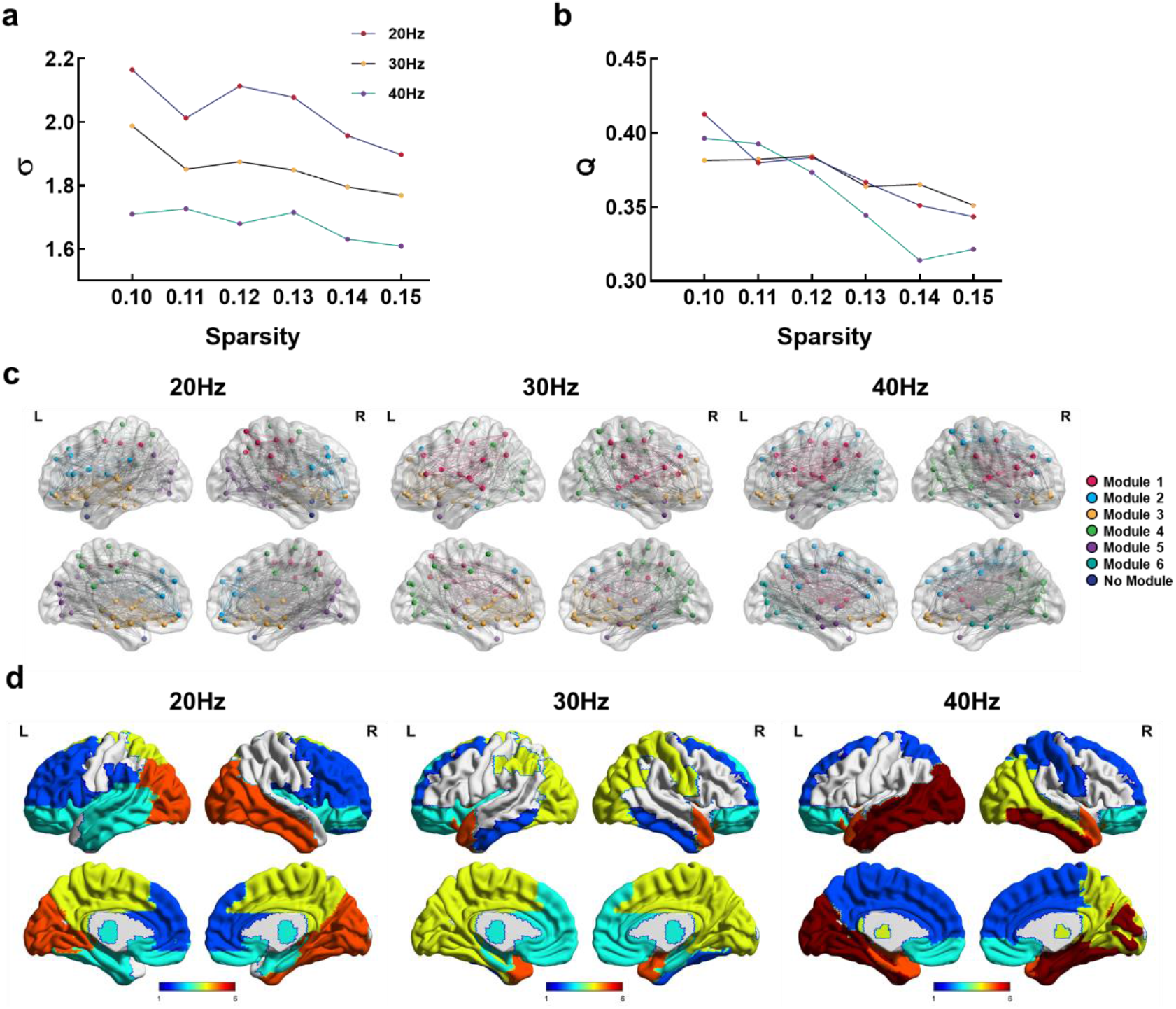
(a) A comparison of the small-world properties of the stiffness-based covariance network in terms of *σ* for the 3 frequencies (0.1≤Sparsity≤0.15). (b) The modularity value Q vs. sparsity for each frequency. (c) Visualization of the stiffness connections in anatomical space (sparsity=0.13). The nodes represented the brain region coordination (AAL90 in MNI) and the edges represented the brain connections. (d) The partitioned modules were mapped in the atlas.

The modularity of the stiffness network decreased with sparsity and the binary biomechanical connection matrices were visualized in 3D anatomical space by a sparsity threshold of 13%(*50*) (**Figure 5**). With sparsity ranged between 0.1 and 0.15, the modularity value Q were all above 0.3. The modular number was 5, 5, and 6 for 20, 30, and 40Hz at a sparsity of 0.13, respectively. By mapping the partitioned modules to the AAL90 atlas, we observed the positions occupied by the partitions were roughly the same as those of front, temporary and occipital lobes, but most of the parietal lobes were not included. With the increase of frequency, the regions out of partitions expand to frontal and occipital lobes. The detailed modular partition results are shown in **Supplement Information Table S4**. The decreased area occupied by the modules also indicated that as the frequency increases, the modulus-based connections were disrupted.

Due to the limit of bearable scan time (limited to ~90 minutes for this study), only 3 frequencies were investigated, and no assessment of the behavioral and cognitive changes of each subject was carried out. Future studies will include patients with mild TBI or depression.

In conclusion, it was observed that vibration to the human brain caused increasing reduction of CBF with increasing vibration frequency, and the regions in which vibration modulated CBF largely overlapped with those responsible for functional connectivity. In addition, increased vibration frequency tended to disrupt the connections of the covariance network based on tissue stiffness. The modulation effect with respect to CBF and stiffness-based network showed an interweaved relationship between the blood flow, stiffness, and the brain functions. Results provided a new perspective for understanding the functional influences of mechanical stimuli to the human brain. Moreover, it showed the potentials of using mechanical vibration as a stimulus to modulate brain in studies of various vibration-related brain diseases.

## 4. Experimental Section

### 4.1 Subjects and ethical approval

52 healthy volunteers (age 25.87±2.78 years old, 31F/21M) participated in this study. All subjects provided informed consent, as approved by the Science and Technology Ethics Committees of Shanghai Jiao Tong University (E2021119I). Among all the volunteers, 7 subjects’ data were excluded due to head motion in scanning, resulting in a total of 45 subjects (26.05±3.09 years old, 26F/19M) for CBF analyses. A total of 41 were finally used for MRE analysis by checking the coefficient of variance (<10%) between two repeated MRE scans.

### 4.2 Study design

To investigate the CBF changes after vibration, we carried out imaging in two sessions (Figure 1): a reference session without vibration (session 1) and with vibration (session 2). Session 1 included a high-resolution T1-weighted (T1W) imaging sequence for anatomical MRI (aMRI), followed by a 3D pseudo-continuous arterial spin-labeling (pCASL) sequence to measure resting CBF (C_0_). At the end of session 1, subjects would rest for 5 minutes. Session 2 included 3 similar imaging modules corresponding to imaging at 3 vibration frequencies (20Hz, 30Hz, and 40Hz). The order of the measurement frequency for each module was carried out in a randomized, counterbalanced way. The vibration motion was induced using a high-frequency accuracy electromagnetic actuator (*38*).

At each imaging module of Session 2, wave images from vibration were also acquired using an MRE sequence. After vibration, CBF was measured immediately using a pCASL sequence. To ensure the reliability and repeatability of the results, the above two imaging steps were repeated for each module. The total time for both sessions was 90 minutes. Subjects were requested with their eyes open during the experiment.

### 4.3 Imaging protocols

All images were acquired on a 3T MRI scanner (uMR790, United Imaging Healthcare, Shanghai, China) with a 24-channel head coil. High-resolution T1W images were acquired to cover the whole brain with a 3D gradient-echo sequence with the following parameters: TR/TE=1068.1/3.4ms, slices=320, FOV=256mm×240mm, voxel size=0.8×0.8×0.8mm^3^. CBF values were measured using a 3D pCASL sequence with the following parameters: 12 tag-control image pairs, 34 transversal slices, TR/TE=4702/14.14ms, label duration=1.8s, post-labeling delay (PLD)=1.8s, FOV=224mm×224mm, voxel size=3.5×3.5×4mm^3^. The CBF values were obtained by:

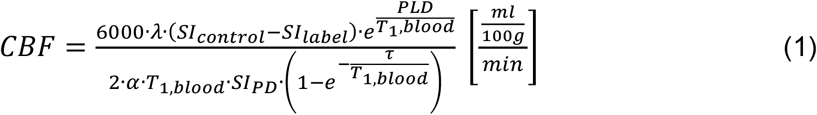

where the brain-blood water partition coefficient *λ* = 0.9 *mL/g*, *SI_control_*, *SI_label_* and *SI_PD_* are the control, label, and proton density-weighted images, respectively, *T*_1,*blood*_ = 1650 *ms* is the longitudinal relaxation time of the blood at 3T, *α* = 0.85 is the labeling efficiency for pCASL, *τ* is the label duration.

MRE was performed during each vibration session (*38*). Single-shot echo-planar imaging (EPI) based MRE sequence with a first-nulling motion encoding gradient (MEG) was used for brain displacement measurement. The MEG had a strength of 30 mT/m and the repetition/time of echo (TR/TE) was 3400/71.6 ms. A total of 34 slices with a thickness of 4 mm was collected for each object. The FOV was 224 mm×224 mm with a matrix size of 128×128. the principal component of the displacement fields was filtered by a 2D Gaussian filter with a kernel size of 5×5 pixels with a standard deviation of 0.65. A fast 3D phase-unwrapping algorithm (*51*) was applied and the stiffness maps were estimated by using a local frequency estimation (LFE) method (*52*).

### 4.4 Data analyses

First, skulls were excluded from T1W images using BET2 (*53*) from the FMRIB Software Library (*54–56*) (FSL v6.0, Oxford Centre for Functional MRI of the Brain, Oxford, UK). Then, CBF and stiffness maps were realigned to T1W images using SPM12 (Wellcome Trust Centre for Neuroimaging, London, UK,). CBF and stiffness maps were resampled into 2 mm^3^ isotropic voxels and normalized to the anatomical standard space defined by the Montreal Neurological Institute (MNI) using Advanced Normalization Tools (ANTs) (*57*). Masks from the automated anatomical labeling atlas (AAL) (*58*) were also used. Each voxel of the CBF and stiffness maps was standardized by dividing the mean CBF/stiffness of the whole-brain for reducing individual variance. Finally, images were smoothed with a Gaussian kernel of 8×8×8 mm^3^ full width at half maximum (FWHM) by SPM12.

Significance analyses were carried out concerning the normalized CBF and stiffness maps using paired student t-tests, two-sample t-tests, or Pearson’s correlation. A two-tailed permutation test with 5000 permutations was used to find significant clusters with the tool Permutation Analysis of Linear Models (PALM) (*59*), and multiple comparisons were corrected by the family-wise error rate (FWER) method with threshold-free cluster enhancement (TFCE) (*60*). A cluster-based threshold of Z > 2.3 was applied. The significance level was set to p-value < 0.05 for the suprathreshold clusters. The cluster analyses were thresholded at t>1.96 or t<-1.96. Toolbox for Data Processing & Analysis of Brain Imaging (DPABI_V6.0 toolkit (*61*),) and MATLAB (MathWorks, Natick, MA, United States) was used for all statistical analysis. Then, the number of overlapped significant voxels in each brain network (MNI space) was estimated for each frequency. The percentage of overlapped significant clusters in all networks and all significant regions were also calculated respectively. In addition, paired student t-tests were made to compare the mean CBF values at the resting state and each of the vibration state at each network for all subjects. Finally, as a prospective study, we calculated the Pearson’s correlations between CBF and stiffness voxel by voxel in MNI space.

### 4.5 Construction of Biomechanical Covariance Matrix

Frequency-dependent correlations of cortical stiffness among different brain regions were analyzed. The cerebral cortex was segmented into 90 regions based on the automated anatomical labelling atlas (AAL) (*58*). Two brain regions were considered biomechanically connected if there is a significant correlation between the average cortical stiffness of the regions across all subjects. A matrix (M×N, where M is the number of subjects of each group, N is the number of brain cortical regions, here N=90) was constructed based on tissue stiffness. Linear regression was performed to remove the effects of age, gender and age-gender interaction for each group. Biomechanical connections were defined as the statistical associations of tissue stiffness between 90 brain cortical regions (45 for each hemisphere) across all subjects. An interregional correlation matrix (90×90) of such connections was acquired by calculating the Pearson’s correlation coefficients across all subjects. Self-connections and negative correlation coefficients were excluded and set to zeros. Multiple comparisons at q<0.05 were corrected by the false discovery rate (FDR) method (*62*) to obtain an interregional stiffness connection matrix *C_ij_* (*i*, *j*: 1, 2…*N*, where *N* = 90).

### 4.6 Graph Theoretical Analysis

The biomechanical connection matrix can be presented by a binarized (0 or 1) and undirected graph (network) G with N nodes and K edges, where nodes and edges indicate brain cortical regions and undirected connections corresponding to nonzero elements of matrix, respectively. By analyzing the small-world properties of G for each frequency, we can investigate the frequency-dependent changes of the biomechanical co-variance network.

In addition, to control the effect that two groups’ graph have different number of edges when using the same correlation threshold, sparsity (S, defined as the existing number of edges in a graph divided by the maximum possible number of edges) was employed as threshold metrics. To estimate small-world parameters properly and minimize the spurious edges, a wide range of sparsity (0.1≤S≤0.15, step 0.01) was used to analyze the biomechanical network properties. As a function of sparsity, discretely varying small-world parameters could also indicate the difference between two groups. All the graph theoretical analysis was performed using Graph Theoretical Network Analysis Toolbox (GRETNA, v2.0.0) (*63*).

### 4.7 Small-world Analysis

Small-world properties (*48*) include two important metrics: clustering coefficient *C_p_* and characteristic path length *L_p_*,which evaluate the extent of interconnectivity of a network at regional and global levels, respectively. Concretely, the clustering coefficient *C_i_* of a node (*i*) is defined as the number of existing connections between the node’s neighbors divided by all their possible connections and *C_p_* is the average of the clustering coefficient over all nodes (*48, 64*):

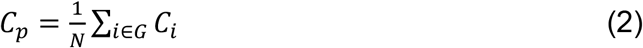

where *N* is the total number of nodes, *G* is the node set in the graph, *C_i_* is the clustering coefficient. Clearly, 0 < *C_p_* ≤ 1, *C_p_* = 1 if and only if the network is fully connected. The characteristic path length *L_ijmin_* of a node is defined as the mean of the shortest path lengths between this node and all other nodes and *L_p_* is the average of all *L_ijmin_*:

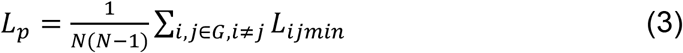

where *N* is the number of nodes, *i* and *j* are node’s index, *G* is the node set in the graph. If a network matches the following criteria, this network would be considered small-world: (1) 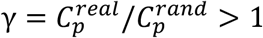; (2) 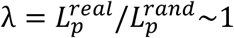 (*48*) or summarized equation σ = γ/λ > 1 (*65*), where the 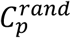 and 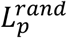 are the mean clustering coefficient and characteristic path length of matched random networks which preserves the same number of nodes and edges and the same degree distribution as real networks (*66*). Here, 100 random networks were generated to compare with real networks. The parameter *σ* of three frequencies were calculated and compared.

### 4.8 Modular Analysis

Using modular analysis(*67*), the brain network was divided into several modules with different self-similarities. For binary undirected network, the modularity Q is defined as(*68*):

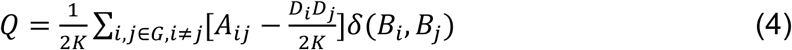

where *K* is the number of exist edges, *i* and *j* are node’s index, *G* is the node set in the graph, *B_i_* is the module to which node *i* is assigned, *δ* is the Kronecker delta function, *D_i_* is the sum of *A_ij_* attached to ties belonging to node *i*.*δ*(*B_i_*, *B_j_*) is 1 if *B_i_* = *B_j_* (and 0 otherwise), *A_ij_*=1 if node *i* and node *j* are connected (and 0 otherwise). A greedy algorithm(*69*) was adopted to maximize Q and detect the modular architecture of the biomechanical co-variance network. All the modular analysis was performed using GRETNA. The binary biomechanical connection matrices were visualized in 3D anatomical space. Each node represented the coordination of one of the ROIs based on AAL, and each line represented two regions that were biomechanical connected. Among all the connections (90×89=8010 pairs of AAL90 regions), the number of connections after FDR correction were 4406 (55.01%),2710 (33.83%), 4108 (51.29%) for 20, 30, and 40Hz, respectively.

## Data Availability Statement

The original contributions presented in the study are included in the article/supplementary material, further inquiries can be directed to the corresponding author(s).

## Ethics Statement

The studies involving human participants were reviewed and approved by the Science and Technology Ethics Committees of Shanghai Jiao Tong University. The patients/participants provided their written informed consent to participate in this study.

## Author Contributions

YF and LK and designed the study. LK and SQ collected MRI data. LK analyzed the CBF data and wrote the protocol. SQ designed the actuator and analyzed the MRE data. LK, SQ, YC, ZH, PH, QH, RZ, XF, LD, YL, FY, GY and YF assisted with the organization of results and interpretation of findings. LK, RZ, XF, LD, GY, and YF wrote and revised the manuscript. All authors have read and approved the final manuscript.

## 5. Acknowledgements

Funding support from the National Natural Science Foundation of China (grant 31870941) and Natural Science Foundation of Shanghai (22ZR1429600) is acknowledged. We would like to thank all our subjects for participating in this research. We thank Prof. Philip V. Bayly, Prof. Guy M. Genin, and Prof. Joshua Shimony from Washington University in St. Louis, and Prof. Mian Long from Institute of Mechanics of Chinese Academy of Sciences for helpful discussions.

## Supplementary Materials for

### Supplementary Text

#### Distributions of the average CBF values at 7 networks

The average CBF values at each network were calculated for all subjects and each vibration. Paired student t-tests showed no significant differences in each network between conditions **(Supplementary Figure S1)**.

#### Distributions of the average CBF values at subregions in DMN

Besides the whole-brain analysis, we also investigate and compare the CBF values at each subregion in DMN **(Supplementary Figure S2)**. We observed significant differences between the vibration mode and the resting state at ACG in DMN. In addition, significant differences were also observed between 20 and 40 Hz vibration and the resting state at the PCC region.

#### A computational model showing correlations between blood flow and tissue stiffness

To investigate the correlation between CBF and stiffness, we constructed a 2D representative volume element (RVE) with a size of 1010×1010 μm^2^ for the brain tissue using COMSOL (**Supplementary Figure S3a)**. The model consisted of two parts: brain parenchyma matrix and vascular system. The vascular system was idealized as tubes uniformly distributed in the RVE. The radius of the tubes was set to be 10um. The brain matrix was taken as a linear solid material with Young’s modulus of 9kPa and Passion ratio of 0.45. The blood was treated as incompressible Newtonian fluid with a viscosity of 2.8×10^-3^ *Pa* · *s*. To study the influence of CBF on mechanical response, the flow speed was set up to vary from 0mm/s to 0.8mm/s with an increment of 0.2mm/s. The apparent shear modulus of the RVE were estimated using uniform simple shear test. The upper and the lower surface of the RVE were displaced 10μm along the x-axis. Periodical boundary condition was set up on the surface of the RVE to ensure uniform deformation. The shear stiffness (G) and CBF was estimated 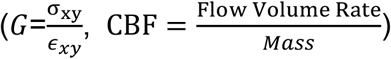 **(Supplementary Figure S3b)**.

#### List of abbreviations of the cortical regions

The cortical surface regions of interest (ROIs) defined by AAL90 template in MNI space was presented **(Supplementary Table S1)**.

#### Regions of decreased CBF

The peak MNI coordinate, peak intensity (t-statistic), voxel numbers of the CBF significantly decreased regions after 30Hz and 40Hz vibration compared with the resting state were reported **(Supplementary Table S2)**. We also reported this information of CBF after 40Hz vibration compared with 20Hz vibration.

#### The regions with significant CBF reduction vs. 7 brain networks

To show the details comparing the regions with a significant decrease of CBF and the seven brain networks, a table showing the number and percentage of overlapping voxels is presented **(Supplementary Table S3)**.

#### Modularity analysis of the biomechanical network

A table summarizing the specific regions for each partition from the modularity analysis is presented, where the number represents the modules which each the region was partitioned **(Supplementary Table S4)**.

**Figure S1.**
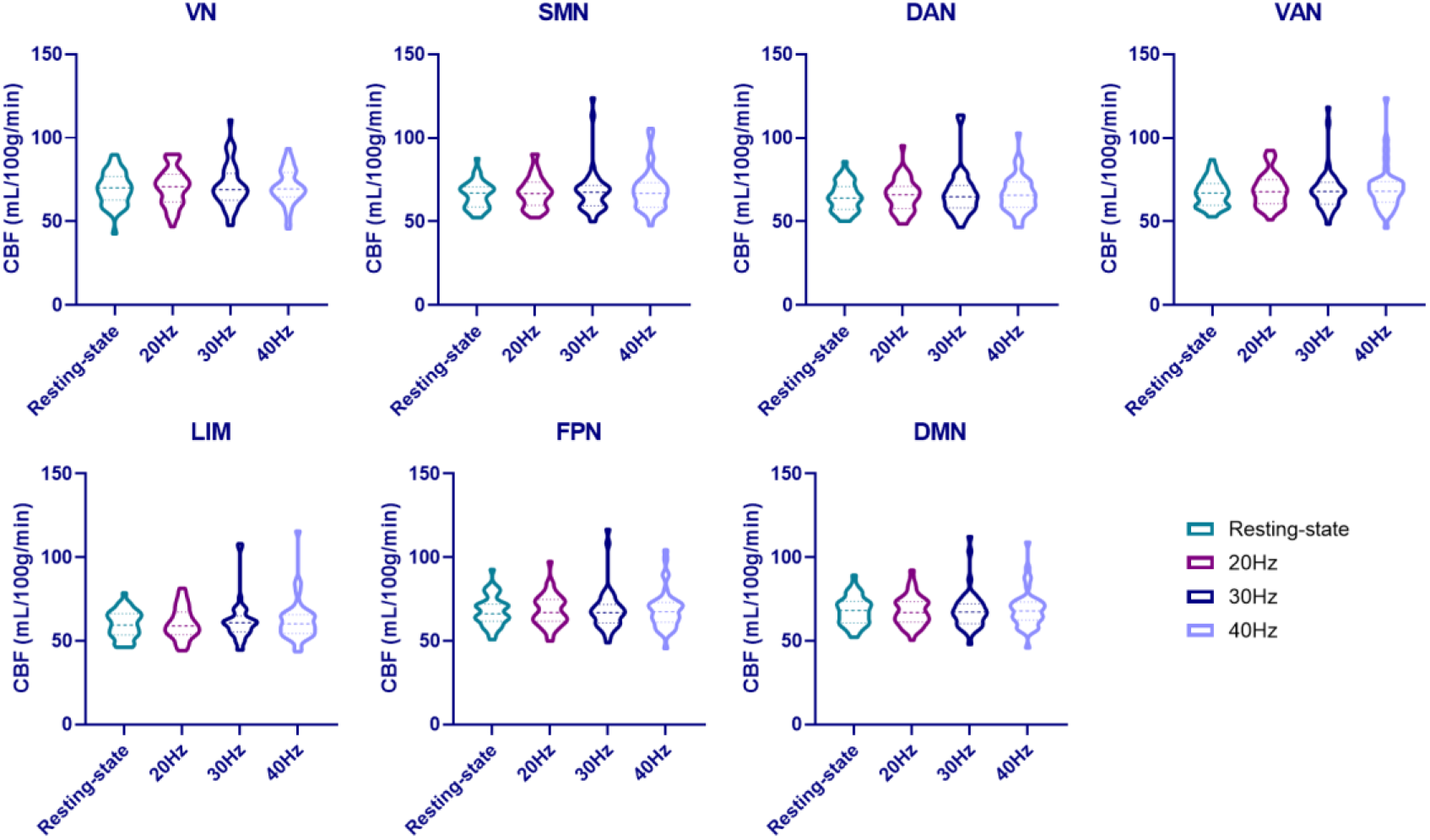
The average CBF in different conditions of 7 networks. There is no significant difference between conditions in each network.

**Figure S2.**
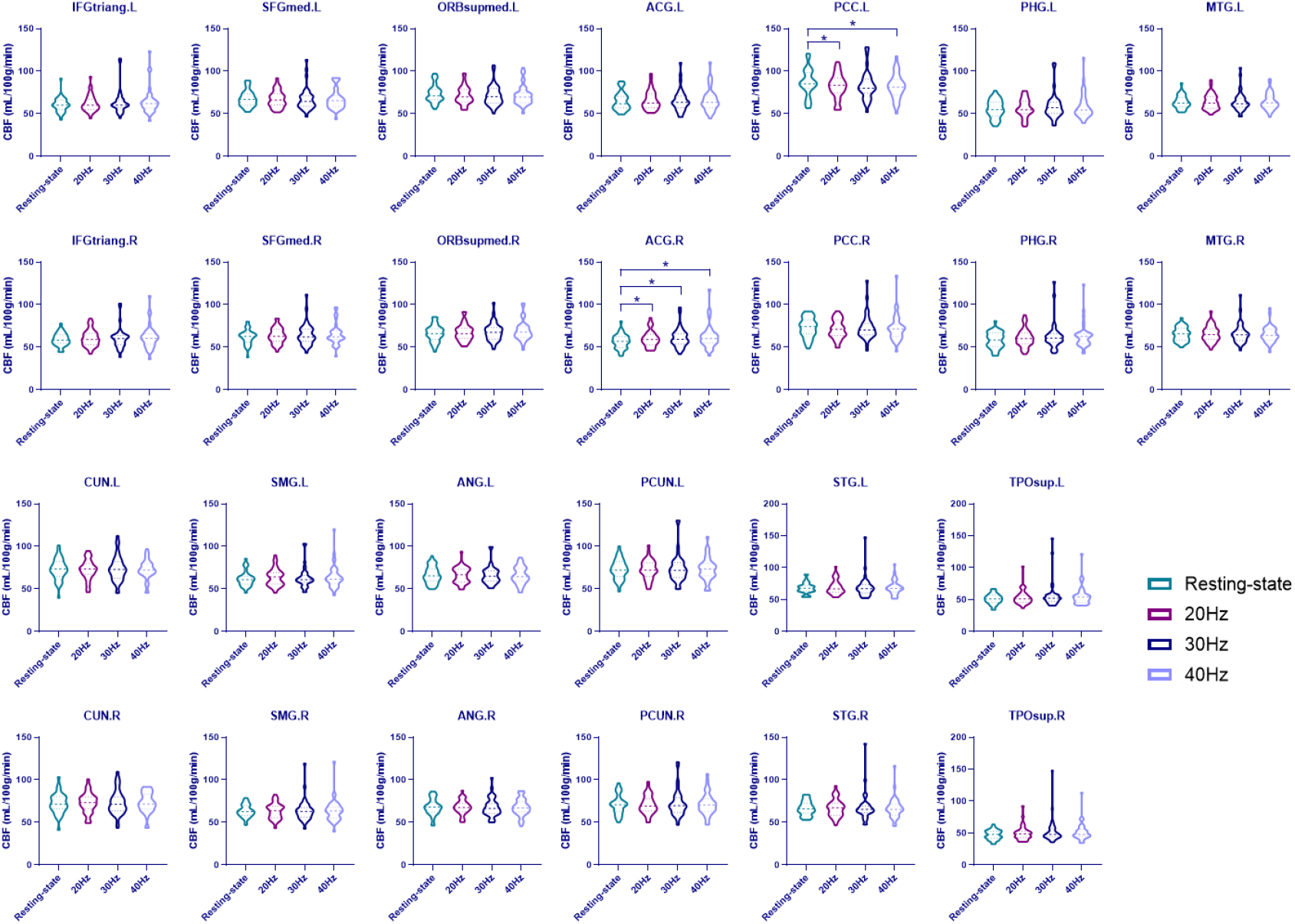
Distributions of the average CBF values at subregions in DMN for the resting state and each vibration frequency. Significant differences were found in left PCC (20/40Hz vs. resting) and right ACG (20/30/40Hz vs. resting)

**Figure S3.**
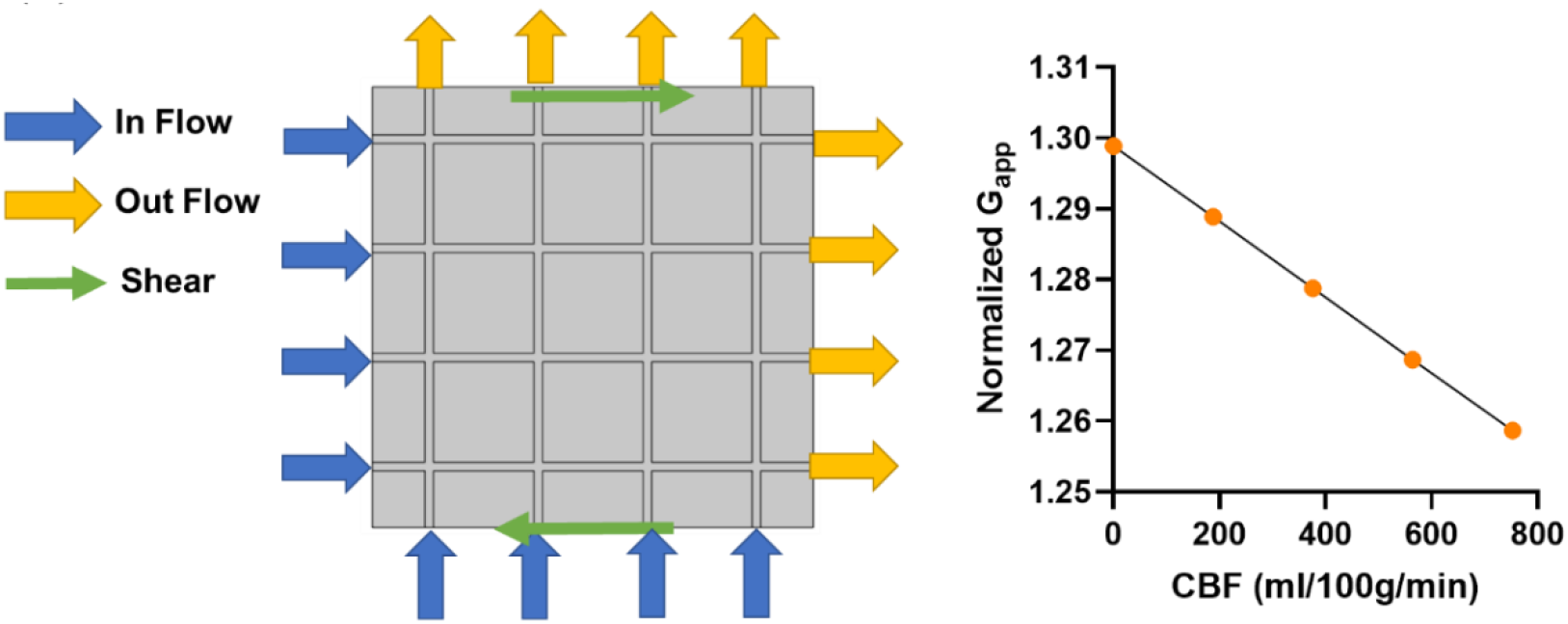
A representative volume element (RVE) based on a fluid-structure interaction (FSI) model constructed using COMSOL. Shear deformation was applied to the RVE to estimate the shear modulus. The normalized apparent shear modulus increased as CBF decreased.

**Table S1.**
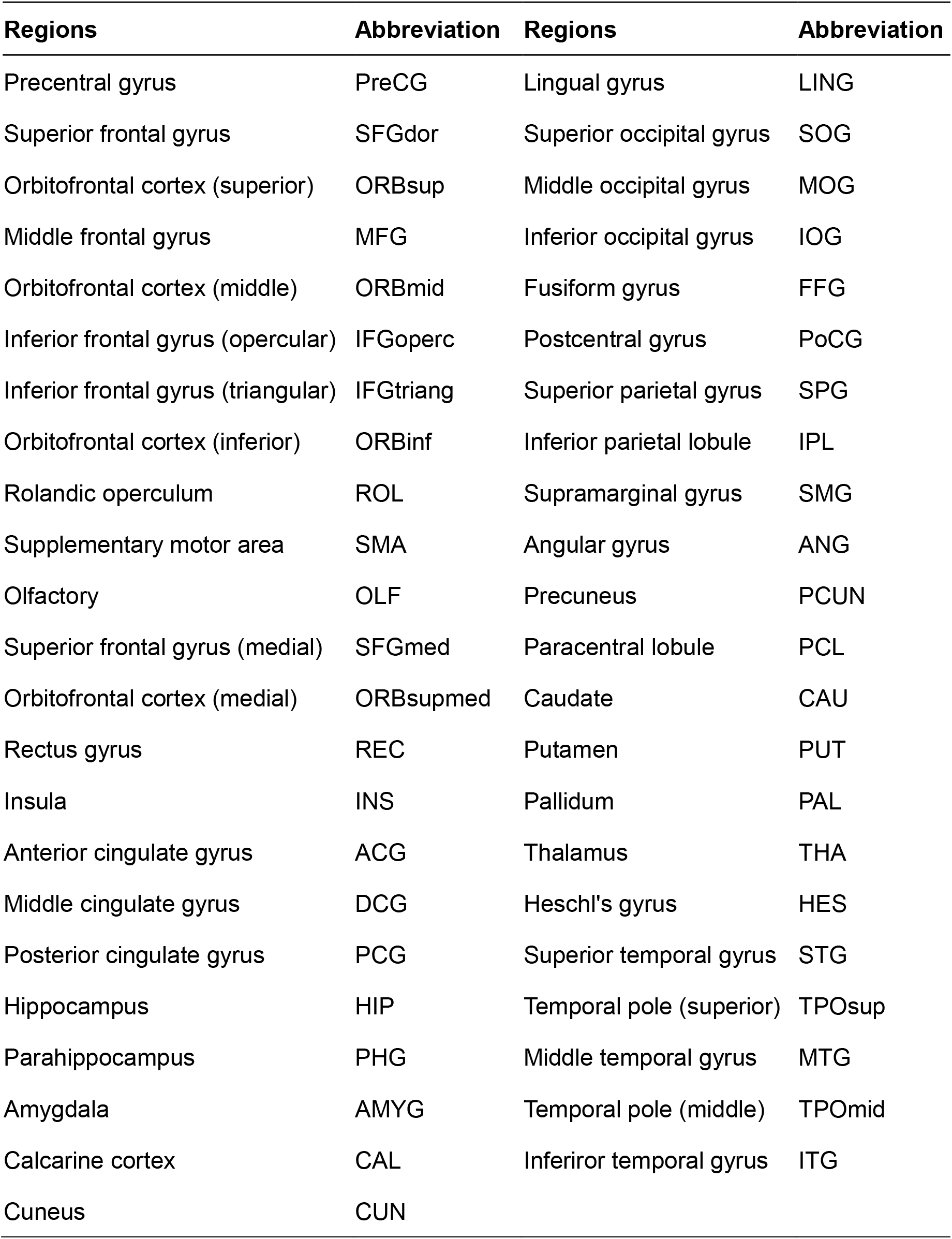
Cortical surface regions of interest (ROIs) defined by AAL90 template in MNI space.

| **Regions** | **Abbreviation** | **Regions** | **Abbreviation** |
| --- | --- | --- | --- |
| Precentral gyrus | PreCG | Lingual gyrus | LING |
| Superior frontal gyrus | SFGdor | Superior occipital gyrus | SOG |
| Orbitofrontal cortex (superior) | ORBsup | Middle occipital gyrus | MOG |
| Middle frontal gyrus | MFG | Inferior occipital gyrus | IOG |
| Orbitofrontal cortex (middle) | ORBmid | Fusiform gyrus | FFG |
| Inferior frontal gyrus (opercular) | IFGoperc | Postcentral gyrus | PoCG |
| Inferior frontal gyrus (triangular) | IFGtriang | Superior parietal gyrus | SPG |
| Orbitofrontal cortex (inferior) | ORBinf | Inferior parietal lobule | IPL |
| Rolandic operculum | ROL | Supramarginal gyrus | SMG |
| Supplementary motor area | SMA | Angular gyrus | ANG |
| Olfactory | OLF | Precuneus | PCUN |
| Superior frontal gyrus (medial) | SFGmed | Paracentral lobule | PCL |
| Orbitofrontal cortex (medial) | ORBsupmed | Caudate | CAU |
| Rectus gyrus | REC | Putamen | PUT |
| Insula | INS | Pallidum | PAL |
| Anterior cingulate gyrus | ACG | Thalamus | THA |
| Middle cingulate gyrus | DCG | Heschl's gyrus | HES |
| Posterior cingulate gyrus | PCG | Superior temporal gyrus | STG |
| Hippocampus | HIP | Temporal pole (superior) | TPOsup |
| Parahippocampus | PHG | Middle temporal gyrus | MTG |
| Amygdala | AMYG | Temporal pole (middle) | TPOmid |
| Calcarine cortex | CAL | Inferior temporal gyrus | ITG |
| Cuneus | CUN |  |  |

**Table S2.** Statistics of the cluster regions with significantly decreased CBF values after vibration.

| Cluster | Number of Voxels | Peak MNI coordinate | Peak MNI coordinate region | Peak intensity (t-statistic) |
| --- | --- | --- | --- | --- |
| **30Hz vs. Resting-state** |  |  |  |  |
| 1 | 672 | [-62, -26, -24] | Temporal_Inf_L | -5.2046 |
| 2 | 1515 | [58, -26, 20] | SupraMarginal_R | -5.2098 |
| 3 | 64 | [-2, 70, 4] | Frontal_Sup_Medial_L | -5.261 |
| 4 | 62 | [-42, -34, 10] | Temporal_Sup_L | -4.6261 |
| 5 | 1962 | [-4, -58, 30] | Precuneus_L | -6.3208 |
| 6 | 146 | [-8, 40, 26] | Frontal_Sup_Medial_L | -4.7671 |
| 7 | 60 | [40, 44, 36] | Frontal_Mid_R | -4.7209 |
| 8 | 288 | [20, 30, 52] | Frontal_Sup_R | -5.0467 |
| 9 | 127 | [44, 6, 52] | Frontal_Mid_R | -5.7009 |
| **40Hz vs. Resting-state** |  |  |  |  |
| 1 | 971 | [-58, -62, -16] | Temporal_Inf_L | -4.7748 |
| 2 | 5032 | [66, -28, 10] | Temporal_Sup_R | -5.9967 |
| 3 | 1212 | [-8, 40, 26] | Frontal_Sup_Medial_L | -4.8526 |
| 4 | 2833 | [30, 52, 32] | Frontal_Mid_R | -4.8465 |
| 5 | 164 | [-62, -40, 14] | Temporal_Sup_L | -3.8191 |
| 6 | 3648 | [-6, -40, 26] | Cingulum_Post_L | -5.1965 |
| 7 | 208 | [42, -14, 46] | Precentral_R | -3.8265 |
| **40Hz vs. 20Hz** |  |  |  |  |
| 1 | 232 | [44, 38, -14] | Frontal_Inf_Orb_R | -5.6062 |
| 2 | 2748 | [68, -28, 10] | Temporal_Sup_R | -5.4656 |
| 3 | 1351 | [40, 42, 28] | Frontal_Mid_R | -4.6984 |

**Table S3.**
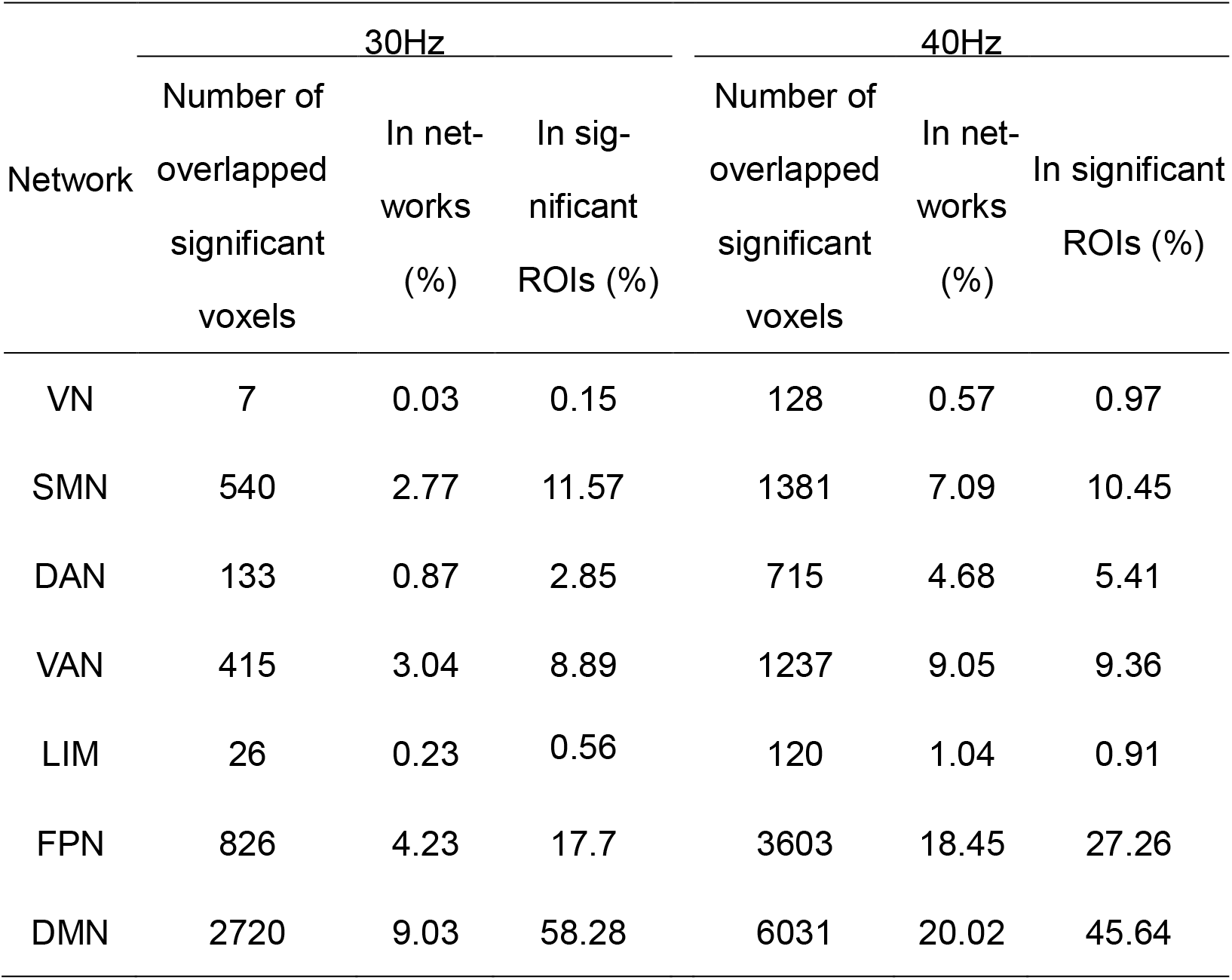
Statistics of the voxels overlapped between the regions with a significant decrease of CBF and the seven brain networks.

| Network | 30Hz |  |  | 40Hz |  |  |
| --- | --- | --- | --- | --- | --- | --- |
|  | Number of overlapped significant voxels | In net-works (%) | In sig-nificant ROIs (%) | Number of overlapped significant voxels | In net-works (%) | In significant ROIs (%) |
| VN | 7 | 0.03 | 0.15 | 128 | 0.57 | 0.97 |
| SMN | 540 | 2.77 | 11.57 | 1381 | 7.09 | 10.45 |
| DAN | 133 | 0.87 | 2.85 | 715 | 4.68 | 5.41 |
| VAN | 415 | 3.04 | 8.89 | 1237 | 9.05 | 9.36 |
| LIM | 26 | 0.23 | 0.56 | 120 | 1.04 | 0.91 |
| FPN | 826 | 4.23 | 17.7 | 3603 | 18.45 | 27.26 |
| DMN | 2720 | 9.03 | 58.28 | 6031 | 20.02 | 45.64 |

**Table S4.** The specific regions for each partition from the modularity analysis. The number represents the modules which each the region was partitioned.

| Regions | 20H<br>z | 30Hz | 40H<br>z | Regions | 20Hz | 30Hz | 40Hz |
| --- | --- | --- | --- | --- | --- | --- | --- |
| PreCG.L | 1 | 1 | 1 | CUN.L | 5 | 4 | 6 |
| PreCG.R | 1 | 1 | 1 | CUN.R | 5 | 4 | 4 |
| SFGdor.L | 2 | 2 | 2 | LING.L | 5 | 4 | 6 |
| SFGdor.R | 2 | 2 | 2 | LING.R | 5 | 4 | 4 |
| ORBsup.L | 3 | 3 | 3 | SOG.L | 5 | 4 | 6 |
| ORBsup.R | 2 | 3 | 3 | SOG.R | 5 | 4 | 4 |
| MFG.L | 2 | 1 | 1 | MOG.L | 5 | 4 | 6 |
| MFG.R | 2 | 1 | 1 | MOG.R | 5 | 4 | 4 |
| ORBmid.L | 3 | 3 | 3 | IOG.L | 5 | 4 | 6 |
| ORBmid.R | 3 | 3 | 3 | IOG.R | 5 | 4 | 4 |
| IFGoperc.L | 2 | 1 | 1 | FFG.L | 3 | 4 | 6 |
| IFGoperc.R | 2 | 1 | 1 | FFG.R | 5 | 2 | 6 |
| IFGtriang.L | 2 | 1 | 1 | PoCG.L | 1 | 1 | 1 |
| IFGtriang.R | 2 | 1 | 1 | PoCG.R | 1 | 4 | 2 |
| ORBinf.L | 3 | 3 | 3 | SPG.L | 4 | 1 | 2 |
| ORBinf.R | 3 | 3 | 3 | SPG.R | 1 | 4 | 2 |
| ROL.L | 2 | 1 | 1 | IPL.L | 1 | 4 | 1 |
| ROL.R | 2 | 1 | 1 | IPL.R | 1 | 1 | 2 |
| SMA.L | 4 | 4 | 2 | SMG.L | 2 | 1 | 1 |
| SMA.R | 4 | 4 | 2 | SMG.R | 1 | 1 | 1 |
| OLF.L | 3 | 3 | 3 | ANG.L | 5 | 1 | 1 |
| OLF.R | 3 | 3 | 3 | ANG.R | 1 | 4 | 4 |
| SFGmed.L | 2 | 3 | 2 | PCUN.L | 4 | 4 | 2 |
| SFGmed.R | 2 | 3 | 2 | PCUN.R | 4 | 4 | 4 |
| ORBsupmed.<br>L | 2 | 3 | 3 | PCL.L | 4 | 4 | 2 |
| ORBsupmed.<br>R | 2 | 3 | 3 | PCL.R | 4 | 4 | 2 |
| REC.L | 3 | 3 | 3 | CAU.L | 3 | 3 | 1 |
| REC.R | 3 | 3 | 3 | CAU.R | 3 | 3 | 1 |
| INS.L | 3 | 3 | 1 | PUT.L | 3 | 3 | 1 |
| INS.R | 3 | 1 | 1 | PUT.R | 3 | 3 | 1 |
| ACG.L | 2 | 3 | 2 | PAL.L | 3 | 3 | 0 |
| ACG.R | 2 | 3 | 2 | PAL.R | 0 | 0 | 0 |
| DCG.L | 4 | 4 | 2 | THA.L | 3 | 3 | 1 |
| DCG.R | 4 | 4 | 2 | THA.R | 3 | 3 | 4 |
| PCG.L | 4 | 4 | 2 | HES.L | 3 | 1 | 1 |
| PCG.R | 4 | 4 | 4 | HES.R | 3 | 1 | 1 |
| HIP.L | 3 | 3 | 5 | STG.L | 3 | 1 | 1 |
| **HIP.R** | **3** | **3** | **6** | **STG.R** | **1** | **1** | **1** |
| **PHG.L** | **3** | **3** | **5** | **TPOsup.L** | **0** | **5** | **5** |
| **PHG.R** | **3** | **3** | **6** | **TPOsup.**<br>**R** | **0** | **5** | **5** |
| **AMYG.L** | **3** | **3** | **5** | **MTG.L** | **3** | **1** | **6** |
| **AMYG.R** | **3** | **5** | **6** | **MTG.R** | **5** | **1** | **4** |
| **CAL.L** | **5** | **4** | **6** | **TPOmid.L** | **0** | **5** | **5** |
| **CAL.R** | **5** | **4** | **4** | **TPOmid.**<br>**R** | **0** | **5** | **5** |
| **ITG.L** | **3** | **2** | **6** | **ITG.R** | **5** | **2** | **6** |

